# Tentative evidence that aging is caused by a small number of interacting processes

**DOI:** 10.1101/2025.04.22.649947

**Authors:** Axel Kowald, Thomas B L Kirkwood

## Abstract

Human life expectancy has increased dramatically over the past two centuries, marking a significant public health achievement. While some projections predict a future where median lifespans reach 100 years, others contend that further longevity will depend on breakthroughs targeting the biological processes of aging. Recent studies in mice have demonstrated that telomerase activation, achieved via gene therapy and transgenic approaches, can extend both median and maximum lifespans substantially without an accompanying increase in cancer risk. We analysed survival data from three such studies using the Gompertz mortality model and show that these interventions reduce the slope parameter, indicative of a slower aging rate, rather than merely lowering baseline mortality. This observation challenges traditional models that assume independent, additive damage accumulation, suggesting instead that aging is driven by a limited number of interdependent processes with significant cross-talk. Mathematical modelling indicates that only three to five processes with substantial cross-talk may account for the observed deceleration. Extrapolation using Swedish survivorship data further implies that a reduction in the aging rate, similar to that seen in mice, could elevate the median human lifespan from 85 to over 100 years. These findings provide a compelling framework for developing targeted anti-aging interventions and a new perspective on the modifiability of the aging process.

## 1. Introduction

Over the past two centuries, human life expectancy has increased dramatically, with no clear evidence of an impending limit to longevity (Oeppen and Vaupel, 2002). This increase, on the order of dozens of years added to average lifespan, stands as a triumph in public health, largely driven by advances in sanitation, nutrition, and medicine. Concurrently, this unprecedented increase in longevity introduces novel challenges, as populations age and societies are confronted with strained healthcare systems and social support networks. In short, the success of humanity in extending life has resulted in the creation of a dual challenge: the support of an ageing demographic.

In contemplating the future, experts diverge in their predictions regarding the potential extent of further increases in human lifespan. Some researchers have used past trends to predict that many individuals born in recent decades will routinely live to 100 years or more (Christensen et al., 2009). Conversely, other demographers and gerontologists posit that the absence of novel interventions targeting the aging process itself renders such extreme longevity improbable. Recent analyses suggest that, under current conditions, the probability of surviving to 100 will remain relatively low, on the order of only a few percent, and that substantial increases in lifespan will require breakthroughs in slowing biological aging (Olshansky et al., 2024).

The diversity of perspectives has given rise to a considerable degree of interest in geroscience, defined as the scientific study of the mechanisms of ageing. Significantly, recent studies in mice have demonstrated that directly modulating an aging mechanism can indeed extend longevity. Specifically, interventions that augment telomerase activity (the enzyme responsible for telomere elongation) have yielded substantial lifespan extension in normal mice. For instance, virus-based gene therapies that activate telomerase in adult mice have been reported to increase median lifespan by approximately 20–40% without obvious adverse effects (Bernardes de Jesus et al., 2012; Jaijyan et al., 2022). In a similar vein, transgenic mice engineered to express high levels of telomerase demonstrate an extension in median and maximum lifespan, accompanied by improved wound healing capacity and a lack of increase in cancer incidence(Zhu et al., 2024). These findings provide compelling proof-of-principle that the aging process can be manipulated in a mammalian model by targeting a single longevity-associated pathway.

We analysed the survivorship curves of these studies by fitting the Gompertz mortality model, which characterises the process of aging by an exponential increase in mortality with age. It was found that treated mice exhibited a reduced Gompertz slope (the parameter defining how rapidly mortality risk accelerates with age), which is equivalent to a reduction of the actuarial aging rate.

From a theoretical standpoint, this outcome is noteworthy. Assuming that the process of aging is caused by several damage accumulating processes, their individual slope parameters should align over evolutionary times. In such a scenario, the elimination of a specific process would be expected to result in a reduction of the Gompertzian intercept parameter, but not the slope parameter. However, the aforementioned mouse experiments challenge this expectation by demonstrating a deceleration of aging (slope reduction), suggesting that the treated animals are not merely living longer, but aging more slowly.

What might account for this surprising deceleration of aging? In this paper, we utilise mathematical modelling to investigate the hypothesis that the biological aging process is governed by a small number of interacting molecular and cellular processes, rather than a large number of independent pathways. The ‘hallmarks of aging’ (Lopez-Otin et al., 2013) consist of several key mechanisms associated with aging. These hallmarks (e.g. telomere attrition, genomic instability, senescent cell accumulation, etc.) are known to be highly interconnected, often engaging in feedback loops and cross-talk. Consequently, an intervention targeting one hallmark may have cascading effects on others. Utilising mathematical modelling and computer simulations, we hypothesise that in mice with activated telomerase, the cross-talk between a limited set of core aging processes contributes to the reduced aging rate. The extension of telomeres has been shown to alleviate or delay other deleterious processes (e.g. genomic instability or cellular senescence), thereby slowing down system-wide aging.

If aging is indeed driven by a small number of interconnected processes, the implications would be far-reaching. The demonstration that the process of aging can be decelerated by targeting a single node of an interdependent network suggests that the process of aging might be more amenable to manipulation than was previously thought. Finally, we show that if these results obtained in mice were transferable to humans, it would correspond to an increase of the median lifespan to around 100 years.

## 2. Telomerase activation reduces the rate of aging

The Gompertz model (Gompertz, 1825) is a mathematical equation that accurately describes the increase of age-related mortality in many species. The model is expressed as follows:

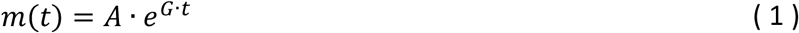

where m(t) is the mortality rate at age t, G is a slope parameter often termed the ‘intrinsic rate of aging’ or ‘actuarial aging rate’, and A is the intercept in a logarithmic plot often termed the ‘basal vulnerability’, which corresponds to the mortality at t=0. In the analysis of survival curves, a linear regression analysis of the logarithm of mortality is a possible approach. However, it should be noted that the calculation of mortality necessitates the binning of data, an approach that is susceptible to errors unless the population size is exceptionally large. An alternative approach that circumvents this issue involves the utilisation of the Gompertz equation to calculate an expression for survivorship, L(t), given by:

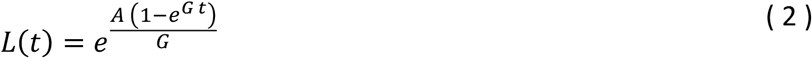

wherein values for A and G are obtained by directly fitting the equation to experimental survivorship data. The NonlinearModelFit function of the Mathematica^®^ software package was employed to estimate the parameters A and G, and it should be noted that this function also provides confidence intervals that can be utilised to ascertain whether there is a statistically significant difference between the parameters of the control and treatment group (see Methods). The Gompertz parameters were fitted to the survival curves of mice from three studies that stimulated telomerase activity either via gene therapy or by creating transgenic strains (Bernardes de Jesus et al., 2012; Jaijyan et al., 2022; Zhu et al., 2024). The survival data, along with the fitted survivorship curves, is displayed in Fig. 1. In all cases, the fitted curves result in r2 values > 0.99, indicating a strong correlation between the model and the data. The subsequent section will provide a detailed description of the three experiments.

**Fig. 1.**
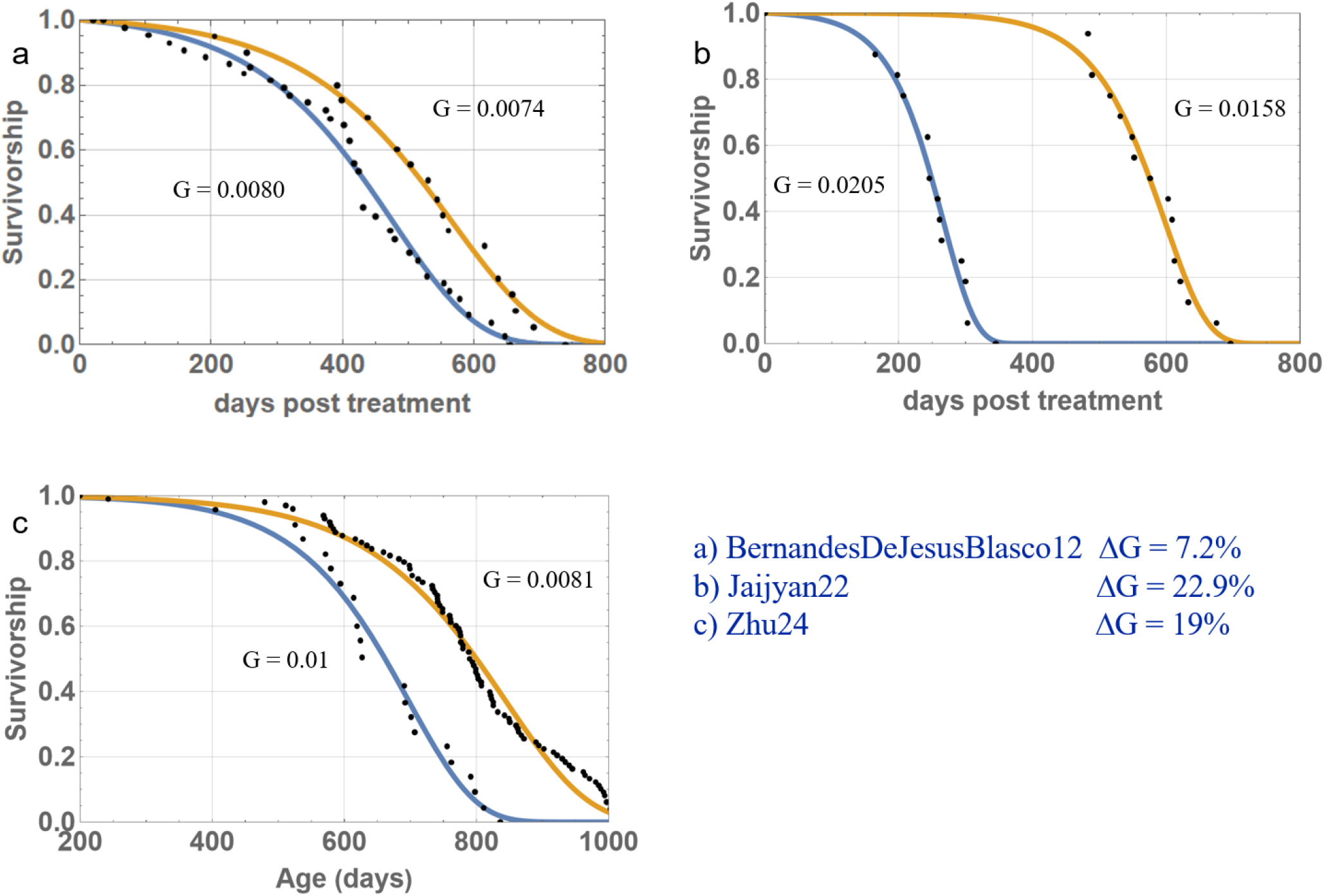
**(a)** Survival data of mice with treated at one year of age with an AAV vector expressing TERT, digitized from Bernardes de Jesus *et al*. (2012), overlaid with survival curves based on fitted Gompertz parameters. Although indicative, the differences in G did not reach statistical significance. **(b)** Survival data for mice with a C57BL/6J background treated at 18 months with a CMV vector expressing TERT, data from Jaijyan *et al*. (2022)), overlaid with survival curves based on fitted Gompertz parameters. Differences in G are statistically significant (p<0.05). **(c)** Pooled survival data of five generations of transgenic mice expressing TERT, digitized from Zhu *et al*. (2024), overlaid with survival curves based on fitted Gompertz parameters. Differences in G are statistically significant (p<0.05). In each case the blue line represents the survivorship curve of controls and the orange line the survivorship curve of the treated group.

In the study by Bernardes de Jesus et al. (2012), adeno associated virus (AAV) was utilised to overexpress the telomerase reverse transcriptase (TERT) in mice with a C57BL6 background, aged one and two years. The median and maximum lifespan (post-treatment) of the treated (n = 21) one-year-old mice increased by 24% and 13%, respectively, compared to the controls (n = 43). The study also demonstrated a concomitant increase in telomere length and a reduction in the prevalence of age-related pathologies and biomarkers. The Gompertz model exhibited an excellent fit (r2=0.996 for the control group and 0.997 for the treatment group, as illustrated in Fig. 1a). The estimates for the slope parameter, G, were 0.008 for the control group and 0.0074 for the treatment group. While this suggests a reduced aging rate in the group with activated telomerase, the difference did not quite reach statistical significance at p=0.05.

Another study utilised a comparable gene therapy approach by employing a cytomegalovirus (CMV) vector to deliver TERT to 18-month-old female C57BL/6J mice (Jaijyan et al., 2022). The vector was administered in monthly intervals, either by intranasal or intraperitoneal injection, and it was observed to increase median and maximum lifespan (from birth) by 40%. For the purpose of our survivorship analysis, survival data from both delivery methods were pooled, resulting in a sample size of 16 for both the control and treatment groups. The Gompertz fit was once again found to be very good (r2=0.993 for the control and treatment group) and resulted in estimated G values of 0.01 (control) and 0.0081 (TERT), which corresponds to a reduction of 22.9% (see Fig. 1b).

Finally, the most recent study that generated transgenic mice of the C57BL/6 strain by inserting a Tert transgene under the control of the elongation factor 1α promoter was examined (Zhu et al., 2024). The authors measured the survival data for five generations of mice and demonstrated that maximum lifespan had been increased by 25%. The study also demonstrated that the mice exhibited enhanced wound healing capabilities and did not demonstrate an elevated cancer risk. For the purposes of analysis, the survivorship data for all five generations was aggregated, yielding a total of 98 mice in the treatment group and 22 in the control group. As in the other cases, the Gompertz fit resulted in a very high correlation coefficient of r2=0.993 for the control group and 0.995 for the treatment group. The slope parameter of the treatment group was 19% lower than the control group, which was statistically significant (p<0.05) (see Fig. 1c).

In summary, the present findings indicate that the activation of telomerase, whether by gene therapy or via the generation of transgenic mice, does indeed result in a slowing of the rise in mortality over time, which is frequently interpreted as the rate of aging.

## 3. A single process should not change the Gompertz slope

Whilst these results are encouraging for research into the ageing process, from a theoretical perspective, it is surprising that the elimination of a single biochemical process involved in the ageing process leads to a reduction in the Gompertzian slope parameter.

The Gompertz law, a phenomenological equation that describes the age-related rise in mortality without reference to any underlying mechanism, can be readily understood and derived if we assume that there is a biological process of damage accumulation, P, driven by positive feedback (see equation (3)). The solution to this differential equation demonstrates that the quantity characterised by this process increases exponentially with time (see equation (4)). If this quantity is also proportional to an organism’s mortality, then the Gompertz law, as shown in equation (1), is recreated.

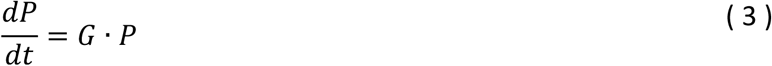

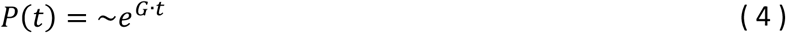

However, it is more probable that there is not a single process of damage accumulation, but several (e.g. the amount of senescent cells, damaged mitochondria, somatic mutations), each contributing some mortality to the overall mortality. Therefore, in the case of three processes, the total mortality rate is expressed as follows:

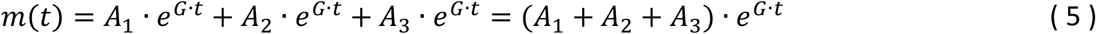

where A_1_, A_2_ and A_3_ represent the process specific intercept parameters and G is a single slope parameter for all processes.

In the field of evolutionary biology, traits frequently exhibit a complex genetic architecture, wherein multiple physiological or molecular processes contribute to a single phenotype (in this case, the rate of aging). However, these different processes rarely contribute equally. One such process may have a larger effect on the trait, whilst others have smaller effects. In the event of one such process demonstrating a substantially larger genetic effect on the rate of aging, and if this process is also accompanied by readily available genetic variation, it is more probable that adaptations will first occur through mutations affecting that major pathway. This phenomenon is formalised as the “line of least resistance” in the field of quantitative genetics(Schluter, 1996; Watanabe, 2024). In the event that an equivalent amount of standing genetic variation is present across all processes, changes in the major pathway are more likely to result in significant phenotypic (and consequently fitness) differences. This differential in the strength of influence that each pathway exerts on the rate of aging leads to a heightened probability of the rapid dissemination of mutations in the major pathway through the population. Consequently, if the slope parameter of a given process is found to be considerably larger than those of the other processes, this process will come to dominate organismic mortality, thus becoming the preferred target for evolutionary adjustments. This dynamic process ultimately leads to a convergence of all slope parameters, resulting in a single value for G in the equation for total mortality. While minor variations might exist, our argument provides a reliable first approximation.

Subsequent examination of equation (5) to ascertain the consequences of the elimination of a solitary damage process leads to the prediction that this will result in a reduction of the overall intercept parameter, but not the slope parameter. This prompts the question of how activation of telomerase (and elimination of telomere-related damage) can reduce the rate of aging.

## 4. Inter-process cross-talk

In the preceding section, it was hypothesised that all processes of damage accumulation are entirely independent of one another. That is to say, it was assumed that physiological changes occurring in one process are completely isolated from the state of the other processes. However, a more realistic scenario is to assume that there is a certain degree of cross-talk between the different processes. For instance, senescent cells might influence the development of cancer, sarcopenia might promote diabetes, and impaired macrophages could foster atherosclerosis. To address such inter-process cross-talk, a set of differential equations was developed to describe the behaviour of the connected processes over time (here exemplified for three processes):

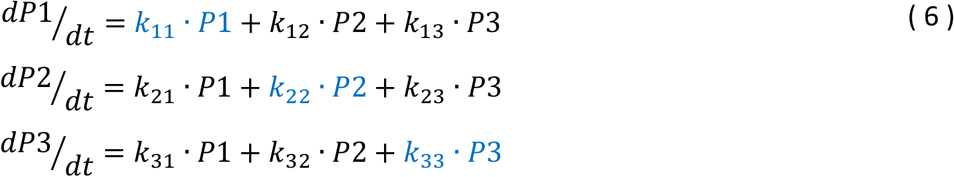

The set of parameters now comprises k_ij_, as opposed to the single parameter G in equation (3). The parameters along the diagonal (k_11_, k_22_, k_33_) control the strength of positive feedback within individual processes, while the remaining parameters govern the cross-talk between processes. As previously discussed, it is hypothesised that k_11_ = k_22_ = k_33_, and for the purposes of simplicity, it is assumed that all cross-talk parameters are identical (but smaller than k_11_, k_22_ and k_33_). The elimination of a single process has now consequences for the accumulation of all other processes. The numerical parameter values for the k_ij_ (and A_i_ to link the processes to mortality) were utilised to reproduce the survivorship curve of Zhu et al. (2024), and the impact of eliminating a single process on the overall aging rate for a specified level of cross-talk was investigated.

The Gompertz slope parameter was calculated by first numerically solving the system of ordinary differential equations (ODEs). The sum of mortalities from the sub-processes was then calculated, and a linear regression was performed on the logarithm of the total mortality (see Methods for details).

As illustrated in Fig. 2, the alteration in the slope parameter, ΔG, is accomplished by the elimination of a single process for a designated level of cross-talk. The magnitude of cross-talk is quantifiable as the ratio of cross-feedback to self-feedback, denoted as k_i≠j_/k_i=j_. Assuming three processes (as is the case in this example), the elimination of one process would require a cross-talk of 33% to achieve a reduction of G of 20%. Conversely, to achieve a 10% reduction in G, a cross-talk of 11.5% would suffice.

**Fig. 2.**
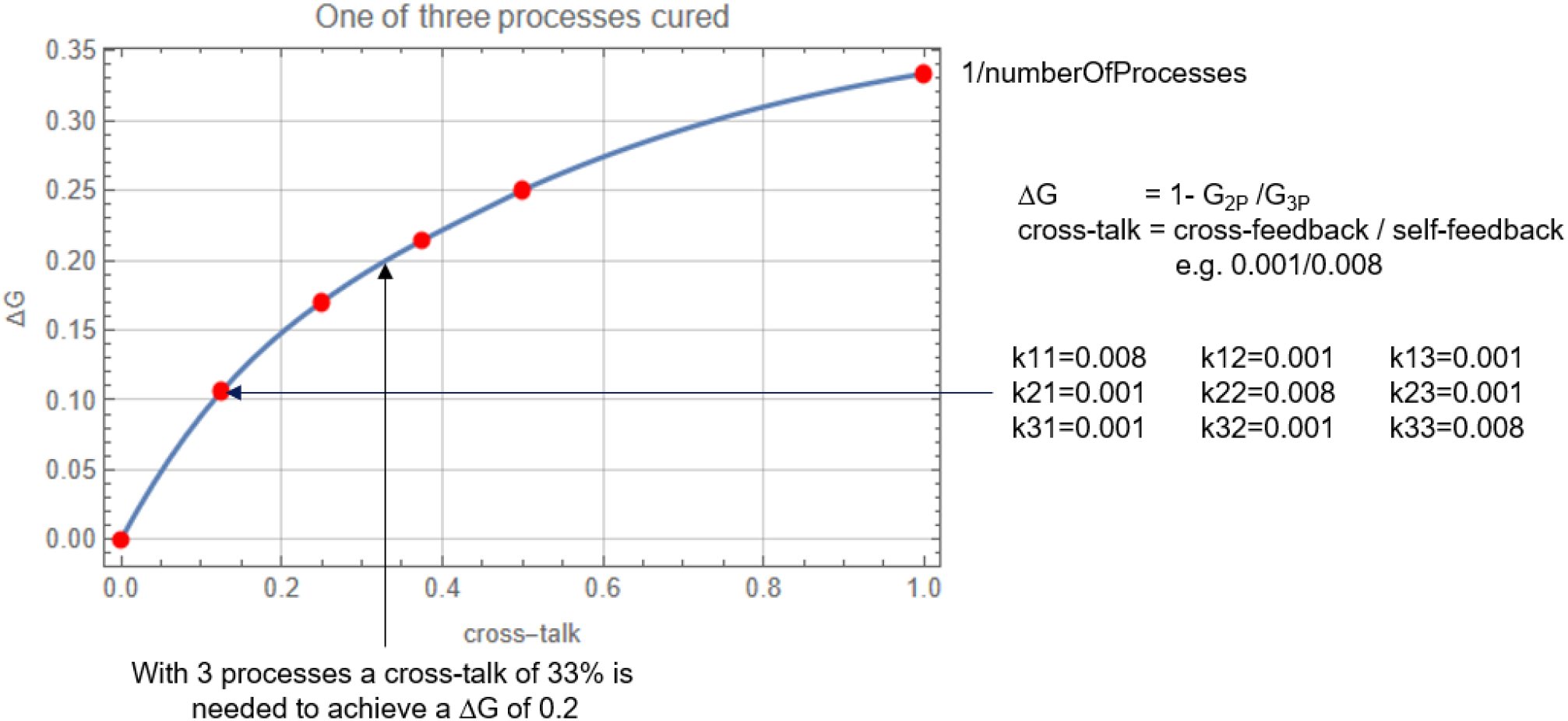
Effect of eliminating one of three processes on the slope parameter for a given level of cross-talk between the processes. Cross-talk is defined as k_i≠j_/k_i=j_ and makes it possible that the elimination of a single process reduces the Gompertz slope parameter up to a maximum degree of 1/numberOfProcesses. For a cross-talk of 0.125 the corresponding set of parameters is shown on the right side.

## 5. How many processes drive mortality?

In the preceding section, it was demonstrated that cross-talk between disparate processes of damage accumulation has the capacity to elucidate how the elimination of a single process can lead to a reduction of the Gompertzian slope parameter, G. The mathematical underpinnings (eqn. (6)) and computations (Fig. 2) were conducted under the assumption of three such processes. However, it must be acknowledged that this number is, in reality, unknown. However, given that experimental reductions of G between 10% and 20% have been observed (Fig. 1), we can ask which level of cross-talk would be required to achieve such a reduction for a given number of processes.

As can be seen in Fig. 3 the number of processes has to be quite small to achieve a 20% reduction through the elimination of a single process. For three processes, a cross-talk of 33% is required (see last section), which increases to 44% for four processes and would reach 100% for five processes.

**Fig. 3.**
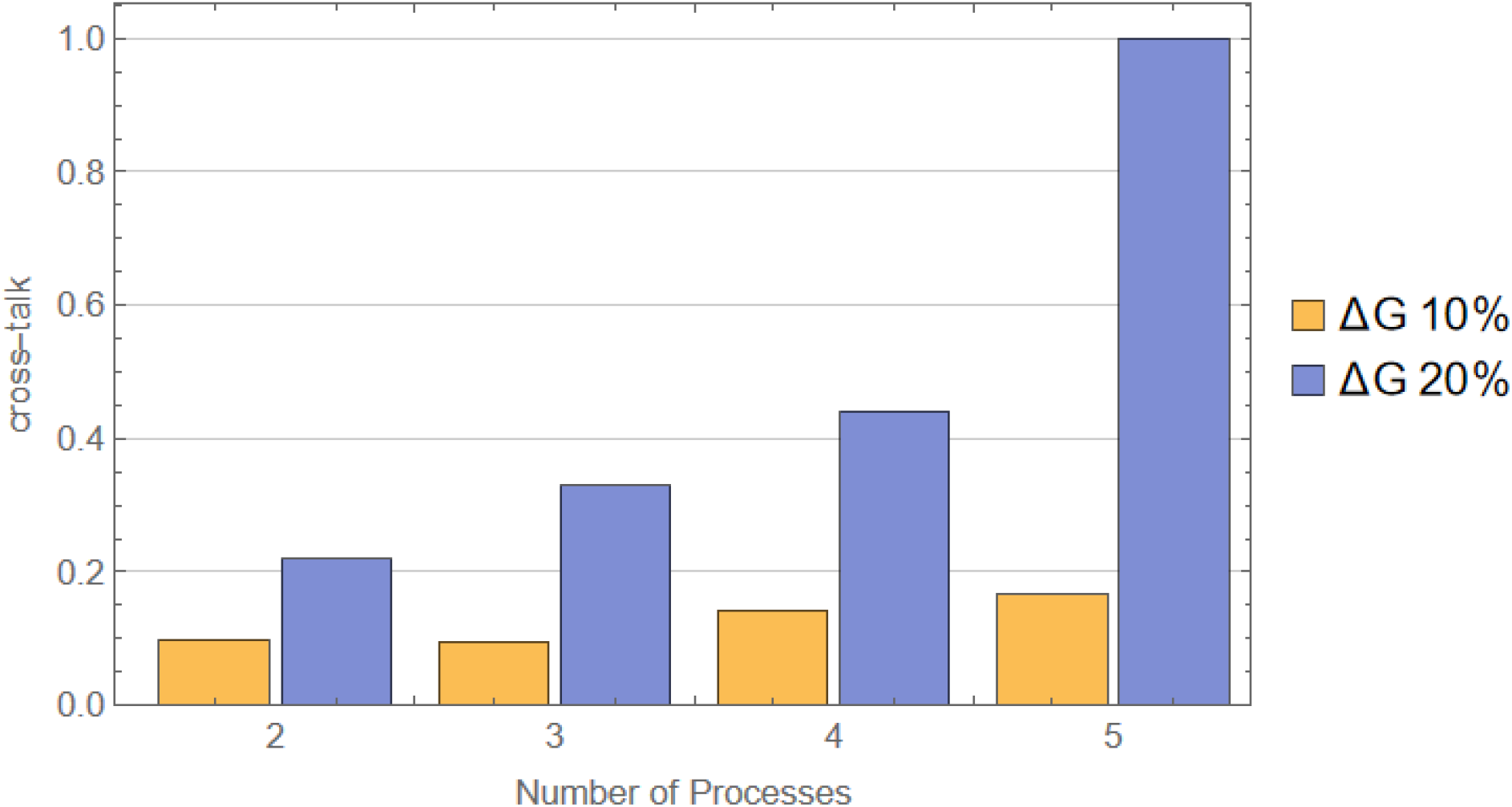
Diagram showing the level of cross-talk that is necessary to reduce the aging rate (i.e. slope parameter) by 10% or 20%, for a given number of damage accumulating processes.

Conversely, to achieve a 10% reduction in G, a lower level of cross-talk is sufficient. However, even in this case, a cross-talk of 100% would be required if there were 10 different processes. Consequently, if experimental findings were to substantiate the hypothesis that telomerase activation can reduce the aging rate by approximately 20%, this would significantly restrict the number of processes that drive mortality.

## 6. Processes with positive feedback

Our analysis of the experimental survivorship data indicates that the aging process may be driven by a small number of processes, which themselves are driven by a positive feedback loop. This finding warrants further consideration, as it is rather unexpected. The prevailing view is that the process of aging is driven by a multitude of factors, with each factor contributing a minimal amount to the overall increase in mortality. A list of nine hallmarks of aging has been proposed (Lopez-Otin et al., 2013), however, over 300 theories of aging have been documented (Medvedev, 1990), many of which focus on molecular damage and functional decline.

However, by focusing exclusively on processes characterised by positive feedback, the actual number of factors may be significantly reduced, as this would eliminate all forms of linear damage accumulation. Furthermore, the identification of which physiological steps are part of the complete feedback loop can be challenging, as the loop can consist of several connected steps. Fig. 4 provides an example of a process that forms a positive feedback loop containing five separate steps (A-E). For the sake of illustration, biological descriptions are assigned to the steps that might lead to a positive feedback, resulting in a continuous progression of atherosclerosis. It is also conceivable that the different steps that form the feedback loop can themselves cause physiological problems and functional decline, which is indicated by X, Y and Z in the diagram. At an organismic level, it would appear that the conditions X, Y and Z increase exponentially with time, but they depend on the central feedback loop formed by steps A to E. Interruption of this feedback loop would also eliminate the rise of X, Y and Z.

**Fig. 4.**
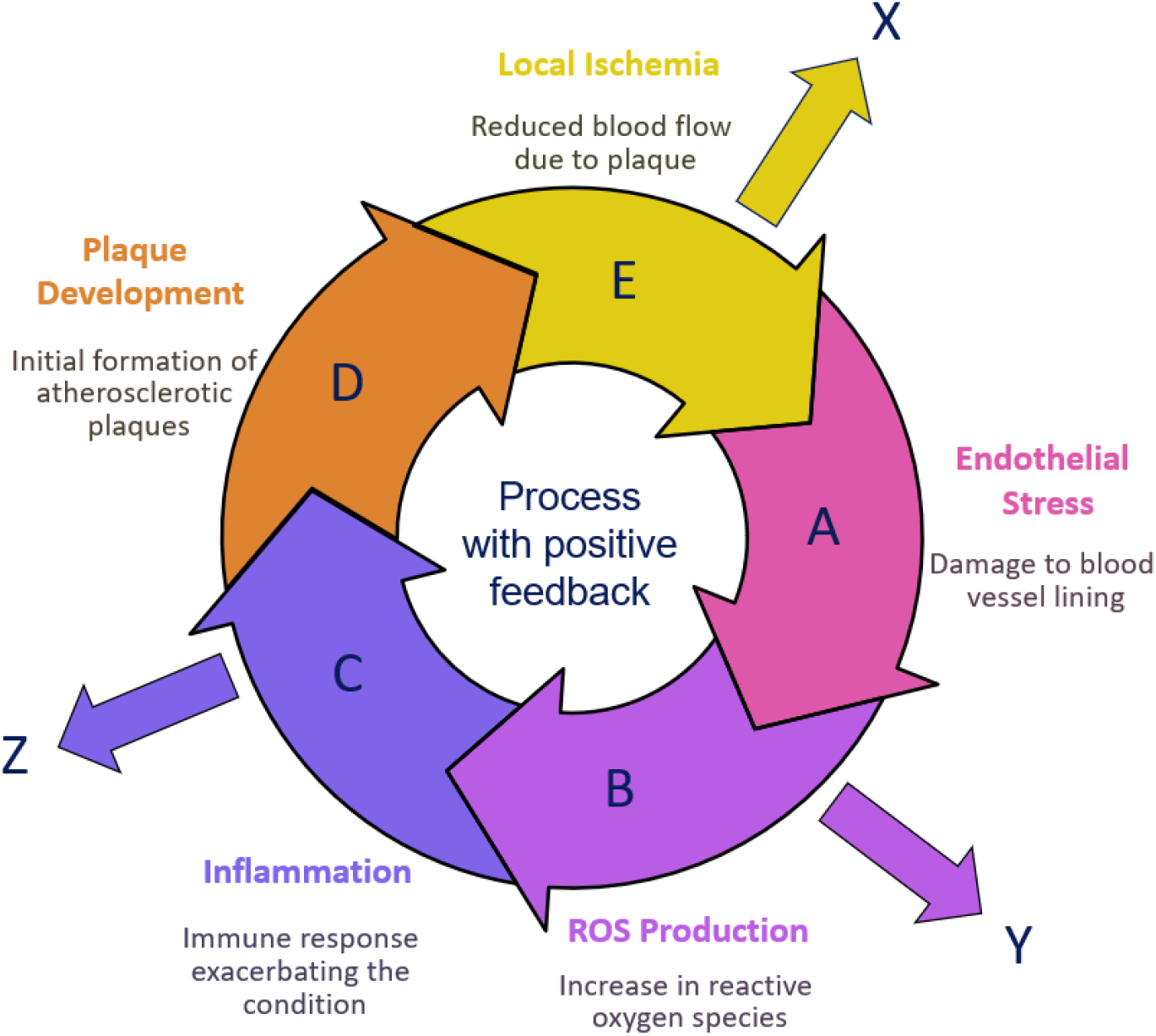
Example of a self-amplifying process consisting of five steps (A-E). For illustration, possible biological events are assigned to the steps such that a positive feedback is generated driving the progression of atherosclerosis. X,Y and Z indicate associated processes that would also grow exponentially but depend on the existence of the central feedback loop. Interrupting or eliminating a single such process that is driven by a positive feedback, might be sufficient to see a substantial drop of the mortality slope.

It is therefore proposed that processes which constitute a functioning feedback loop may be uncommon and could give rise to other conditions that, in isolation, do not form a feedback loop. This could provide a rationale for the substantial decrease in the aging rate observed following the elimination of a single damage process (Bernardes de Jesus et al., 2012; Jaijyan et al., 2022; Zhu et al., 2024).

## 7. Discussion & Conclusions

The objective of ageing research is to decelerate the ageing process and extend the healthy lifespan. Recent experiments that activate telomerase have shown significant reductions in mortality rates, accompanied by increased median and maximum lifespan (Bernardes de Jesus et al., 2012; Jaijyan et al., 2022; Zhu et al., 2024). These findings represent a significant success of this research. Furthermore, the magnitude of the achieved reduction in the aging rate is a significant piece of information that can provide further insight into the structure of the aging process. In the event that the process of aging is attributable to multiple, discrete processes, the elimination of a single process should result in a reduction of the Gompertzian intercept parameter, but not the slope parameter. A plausible hypothesis is that the implicated processes are interconnected via a certain degree of cross-talk, whereby a specific process not only accelerates its own progression, but concurrently (though to a lesser extent) influences the progression of the other processes. This hypothesis is more realistic than the alternative of complete independence, as it suggests a complex interconnection between the processes involved in the ageing process.

The present study has formulated a mathematical framework to facilitate the investigation of the effects of eliminating a single process on the resulting mortality slope. The findings of this study indicate that a cross-talk of 33% is sufficient to explain the 20% reduction in the aging rate observed in certain experiments (Jaijyan et al., 2022; Zhu et al., 2024), under the assumption that a total of three processes are driving aging. However, if a greater number of processes were found to be responsible for the ageing process, a higher degree of cross-talk would be required, with 100% being reached for five processes. A reduction of the mortality slope by a mere 10% (comparable to the results of Bernardes de Jesus et al., (2012)) would necessitate less cross-talk and would be compatible with up to ten processes. In the event that the study of Zhu et al. (2024) is to be regarded as a reference point (due to its superior level of detail and the largest number of animals involved), then the mathematical analysis performed by the present authors would suggest that the process of aging is driven by a mere three or four processes, which are connected via a substantial amount of cross-talk (30%-40%).

This outcome is noteworthy in view of the numerous molecular processes that incur damage over time (e.g. telomere shortening, defective mitochondria, epigenic patterns, etc.) and the many physiological functions that exhibit a decline (e.g. muscle strength, immune system, atherosclerosis, etc.). However, for mortality to increase exponentially, some sort of self-enforcing damage accumulation appears to be necessary, and the number of processes with a positive feedback loop might be rather small. Furthermore, if the individual steps of such a self-amplifying process might themselves cause damage and mortality in other parts of the organism (see Fig. 4), then a small number of feedback driven processes might actually account for a large number of age-related phenotypic changes. This finding is consistent with the observation that telomerase activation not only leads to life extension but also improves a range of other conditions, including wound healing, glucose tolerance, physical performance, and maintenance of body mass (Jaijyan et al., 2022; Zhu et al., 2024).

At present, the number of publications containing good survivorship data after telomerase activation is relatively small. However, if further studies would corroborate these findings, it would have important consequences for research into ageing. The most significant would be the realisation that it is indeed feasible to decelerate the aging rate and extend lifespan by manipulating individual pivotal processes. Although the experiments conducted thus far have only demonstrated a 20% reduction in mortality slope, the potential implications of such findings, if applicable to humans, could be immense. A recent publication has concluded that a radical life extension in humans is very unlikely in the foreseeable future (Olshansky et al., 2024), based on the fact that the increase in life expectancy has actually slowed down over the last 30 years. The authors of the analysis focused on past trends in eight countries and predicted that a median lifespan of 100 years is implausible in this century. In order to investigate the consequences of a 20% reduction in the aging rate on the human population, we analysed Swedish cross-sectional survivorship data from 2022 (Human-Mortality-Database) using the Gompertz model (Fig. 5). The original data demonstrated a median lifespan of 85 years, which increased substantially to 104 years when the slope parameter was reduced by 20%. Consequently, the present experimental methodologies have the potential to effect a radical life extension in humans, provided that the animal data can be extrapolated.

**Fig. 5.**
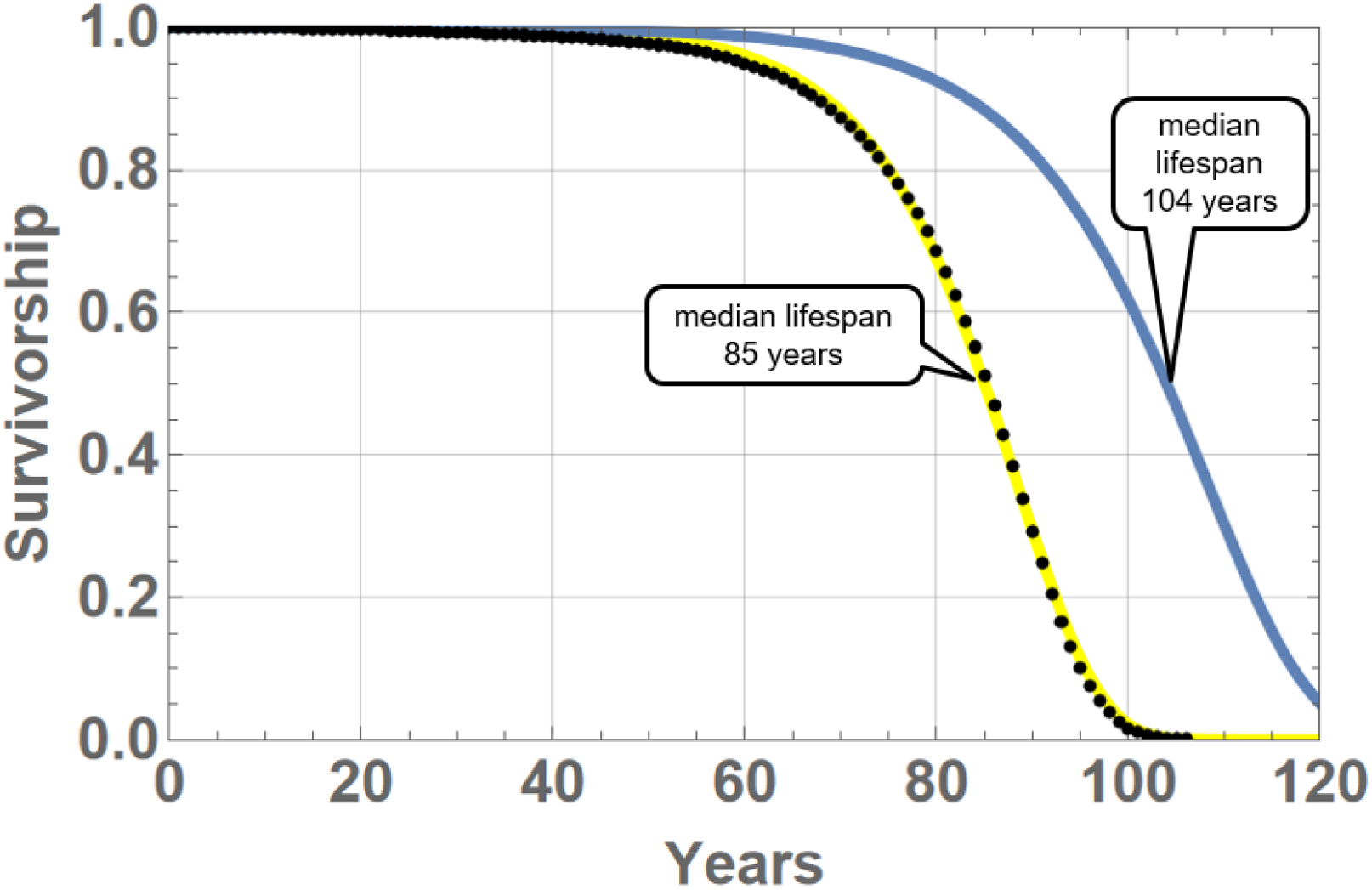
Swedish survivorship data from 2022 (black dots) (Human-Mortality-Database) together with a curve based on a fitted Gompertz model (yellow line). If the slope parameter could be reduced by 20%, the median lifespan would increase to 104 years, quite in contrast to the predictions of (Olshansky et al., 2024).

## Methods/SI

### Data Acquisition

To perform the analysis of survivorship curves for the three studied publications (Bernardes de Jesus et al., 2012; Jaijyan et al., 2022; Zhu et al., 2024) the data are needed as numerical values. Jaijyan et al. (2022) provided survival times in text form with a precision of 0.1 months, while the data from the other two publications were obtained by digitizing their Fig. 3 (Bernardes de Jesus et al., 2012) respectively Fig. 2 (Zhu et al., 2024) using the software LabPlot 2.11.1 (https://labplot.org/).

### Survival Curve Fitting

The survivorship curve of the Gompertz model (eqn. (2)) was fitted to the experimental data using the software package Mathematica (v14). Specifically, the function NonlinearModelFit was used to estimate values for the parameters *A* and *G*. The function also provides confidence intervals which can be used to decide if there is a statistically significant difference between the value of the parameter G of the control and treatment group. First a fit to the control group was performed to obtain an estimate for G_control_. In the following fit to the treatment group the G parameter was then expressed as (G_control_ + deltaG) and NonlinearModelFit was used to obtain an estimate for deltaG. If the confidence intervals of deltaG encompassed zero, this was interpreted as a non-significant difference between both groups.

### Calculating mortality slopes (G) of the multi-process model

We used Mathematica (v14) to numerically solve the coupled set of ordinary differential equations (ODE) given by eqn. (6) for a given set of parameter values k_ij_ and A_i_. Total mortality was then calculated as the sum of mortalities of the sub-processes as described in the main text. Finally, a value for parameter G was obtained by performing a linear regression on the logarithm of the total mortality. The regression was performed over the 25%-75% percentile range of the simulated timespan (i.e. from 250-750 days for a 1000 day simulation). This is necessary since the logarithm of mortality does not follow a perfectly straight line for early ages. The reason is that it is assumed that a ‘cure’ of a process stops its progression (G=0), but does not remove the base value that is already present at young ages (A>0). In addition to the numerical value for the calculated value of the slope parameter G, the routine also produces several diagrams for visualization. Fig. S1 shows an example output for the specified parameters. Please note that here different values were used for k_11_, k_22_ and k_33_ to show all of the resulting curves (otherwise the curves of all sub-processes would be identical and indistinguishable).

Finally note that for the simulation of four or five connected processes, as is necessary for Fig. 3, the system of ODEs was correspondingly modified.

**Fig. S1:**
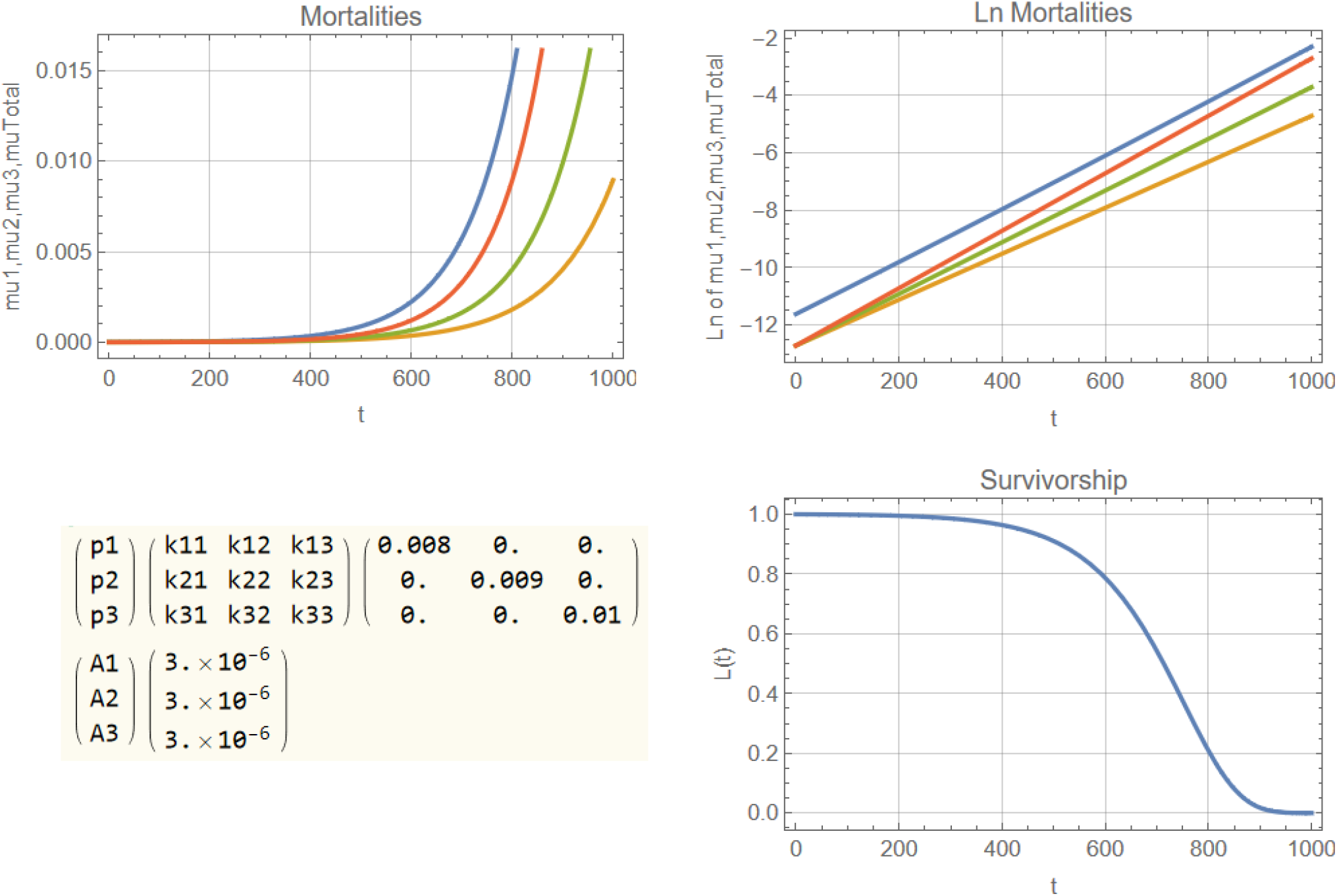
Graphical output of the simulations of the multi-process model (eqn. (6)). For demonstration purposes different values for k_ii_ have been used so that the mortality curves of all three processes can be seen together with the total mortality (blue). The diagram shows the mortality values (top left), the logarithm of the mortalities (top right) and the resulting survivorship curve (bottom right).

